# Oncogenic addiction to high 26S proteasome levels

**DOI:** 10.1101/211300

**Authors:** Peter Tsvetkov, Julia Adler, Nadav Myers, Assaf Biran, Nina Reuven, Yosef Shaul

## Abstract

Proteasomes are large intracellular complexes responsible for the degradation of cellular proteins. The altered protein homeostasis of cancer cells results in increased dependency on proteasome function. There are several different proteasome complexes that may be assembled in cells, with the 20S catalytic core common to them all. 20S proteasomes can function in isolation, or as part of larger complexes (26S) with regulatory particles (RP) such as the 19S that is needed for the targeting and processing of ubiquitinated substrates. Proteasome inhibitors target the catalytic barrel (20S) and thus this inhibition does not allow the deconvolution of the distinct roles of 20S vs. 26S proteasomes in cancer progression. We examined the degree of dependency of cancer cells specifically to the level of the 26S proteasome complex. We found that oncogenic transformation of human and mouse immortalized cells with mutant Ras induced a strong increase in the translation of the 26S proteasome subunits, giving rise to high 26S complex levels. We show that depletion of a single subunit of the 19S RP was sufficient to significantly reduce the 26S proteasome level and lower the cellular 26S/20S ratio. We further demonstrate that the accumulated 26S proteasome was essential for the viability of the transformed cells. Moreover, the viability of 20 different cancer cell lines, but not normal human fibroblasts, was severely compromised upon specific 26S proteasome suppression regardless of their p53 status. Suppression of 26S activated the UPR and Caspase-3, which at least partially explains the cell-killing effect. Morphologically, suppression of the 26S proteasome resulted in cytoplasm shrinkage and nuclear deformation. Thus, the tumor cell-specific addiction to high 26S proteasome levels sets the stage for future strategies in cancer therapy.

## Introduction

Proteasomal protein degradation is crucial in maintaining cellular integrity, in regulating cell cycle and in cell fate determination such as proliferation and cell death (1). Proteasome degradation of proteins is mediated by two distinct proteasome complexes; the 26S and the 20S proteasomes. The 26S proteasome consists of the 20S catalytic domain assembled with either one or two 19S regulatory particles (RP) (2, 3). In the well-characterized ubiquitin-proteasome system (UPS) a protein substrate is targeted to the 26S proteasome following conjugation of a poly-ubiquitin chain *(1, 4)*. The poly-ubiquitinated substrate is then recognized by specific subunits of the 19S RP of the 26S proteasome where it is de-ubiquitinated, unfolded by the ATPases and translocated into the 20S catalytic chamber for degradation (2, 5, 6) Over the years an alternative, ubiquitin-independent proteasome degradation pathway has been described whereby intrinsically disordered proteins (IDPs) such as p53, c-Fos, BimEL (7–9) (and others as reviewed in ***(10, 11)***) are degraded by the 20S proteasome in a process that does not involve active ubiquitin tagging ***(12, 13)***. Thus, there are at least two distinct proteasome protein degradation pathways, each regulated by the distinct 26S and 20S proteasome complexes.

The UPS as a regulator of cell death has been a tempting target for drug development for many pathologies, including cancer (14–16). Various tumors have been shown to express high levels of proteasome subunits and higher proteasome activity (17, 18). A number of studies suggest that cancer cells exhibit high sensitivity to proteasome inhibition (19). Proteasome inhibition is specifically efficient in treating lymphoid malignancies, particularly multiple myeloma where the proteasome inhibitor bortezomib (VELCADE, PS-341) was approved for therapy (20). Proteasome inhibitors were also shown to be efficient in various screens of solid and hematologic tumors (19, 21) and currently there is an increasing number of cancers that are in the process of clinical trials with different proteasome inhibitors (22). Proteasome inhibitors such as bortezomib, MG132, and carfilzomib inhibit the catalytic activity within the 20S proteasome particle that is essentially responsible for the activity of both the 20S and the 26S proteasomes (23). Therefore, these drugs cannot be utilized to characterize or distinguish between any unique functions that either the 20S or the 26S proteasome complex play in the cell.

The cellular levels and 26S/20S proteasome complex ratio is both dynamic and regulated. Recent studies showed that a common mechanism of resistance to proteasome inhibitors involves the reduction in the cellular 26S/20S proteasome complex ratio (24–26). These findings highlight the notion that the ratio of proteasome complexes in the cell is a regulated and crucial process. Further findings demonstrate the functional alteration of the 26S/20S proteasome complex ratio in the context of cell cycle progression (27), neuronal function (28, 29), metabolic regulation (30–33) and aging (34–36).

During the process of transformation there is increased dependency on proteasome function as part of global increased burden on the protein homeostasis machinery (37). Genetic screen analysis in several models including Ras transformation and triple negative breast cancer cells revealed a strong dependency on the proteasome function (38–40). These findings motivated us to explore the alteration in the 26S-20S proteasome complex levels and ratio in oncogenic transformation. Utilizing an isogenic cell line model, we challenged the prediction that cancer cells are specifically vulnerable to the reduction of the 26S proteasome complex. The result of our study led us to the discovery that highly transformed cells exhibit increased levels of the 26S proteasome complexes and are extremely sensitive to their specific suppression.

## Results

### 26S proteasome levels increase following H-Ras V12 transformation

Little is known on the alteration of the different proteasome complexes (26S and 20S) during the process of cancer transformation. Initially we set out to examine the specific levels of the 26S proteasome upon cell transformation. To this end we overexpressed H-Ras V12 (G12V) to transform the immortalized NIH3T3 cells (Figure 1a). The level of the PSMA4, a component of the 20S proteasome and PSMD1, a component of the 19S RC, were partially increased after transformation (Figure 1b). However, in the transformed cells the level of the assembled 26S proteasome, in particular the double caped complex, was markedly increased whereas the level of the 20S was concomitantly decreased (Figure 1c).

**Figure 1:**
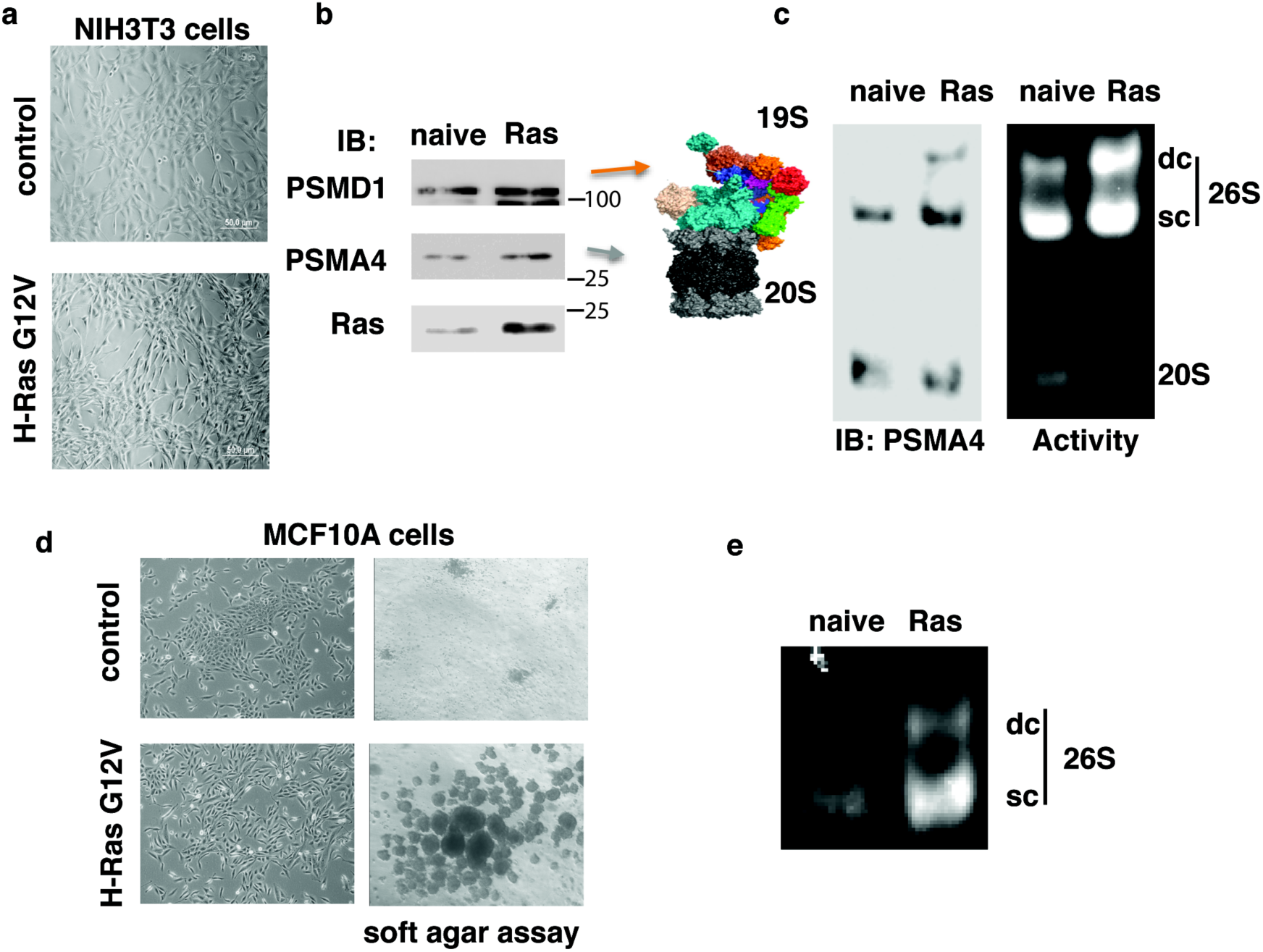
The H-Ras G12V transformed NIH3T3 and MCF10A cells contain high levels of 26S proteasome. **(a)** NIH3T3 cells were transduced with retroviruses carrying pBabe H-Ras G12V gene, and visualized by microscopy. The cells displayed a transformed phenotype with a spindle-shaped and highly refractile morphology. **(b)** Expression of the proteasomal subunits PSMA4, a 20S component, and PSMD1, a 19S component, was quantified by immunoblot (IB). **(c)** 26S and 20S proteasomal complex activity and level were analyzed by native gel electrophoresis in both naïve NIH3T3 and Ras transformed cells. The 26S complex is either double capped, namely each of both ends of the 20S proteasome is occupied by a 19S complex (DC-26S) or single capped (SC-26S). **(d)** MCF10A were transduced with retroviruses carrying pBabe H-Ras G12V gene, visualized by microscopy and subjected to soft agar assay. Transformation was evidenced by a fibroblast-like appearance and a dispersed cell distribution on a regular culture dish (left) and by anchorage-independent growth in soft agar (right). **(e)** 26S proteasomal complex activity was analyzed by native gel electrophoresis in both naïve MCF10A and Ras transformed cells.

Next, we transformed the human immortalized MCF10A cell line. The transformed cells formed colonies in soft agar gel, suggesting they acquired anchorage independent growth, a hallmark of transformation (Figure 1d). In the transformed cells the levels of the assembled and functional 26S proteasome were significantly higher than in the naïve (immortalized) state (Figure 1e). These data suggest that H-Ras V12 transformation of mouse and human cells is associated with an increase of functional 26S proteasome complexes.

### Proteasome subunits accumulate in H-Ras G12V-transformed MCF10A cells

The H-Ras V12-mediated increase in 26S proteasome complex levels could be due to either a transcriptional or post-transcriptional event. No significant differences in the 19S PSMD (Figure 2a) and PSMC (Figure 2b) subunit mRNA levels were observed between the immortalized and transformed cells. As a positive control, we quantified the mRNA levels of CTGF, a hallmark of transformation, which was upregulated in H-Ras V12 transformed MCF10A cells (Figure 2c). However, immunoblot analysis revealed higher protein levels of the selected members of the 19S RC (PSMDs and PSMCs) and the 20S (PSMA1) subunits in the MCF10A transformed cells (Figure 2d triplicates). Interestingly, the obtained fold of increase was similar to that of H-Ras V12 level, despite the fact that the Ras mRNA level was five folds higher in the transformed cells. These data suggest that upon transformation cells accumulate much higher level of the 26S proteasome by improving the translation and/or stability of the different proteasome subunits.

**Figure 2:**
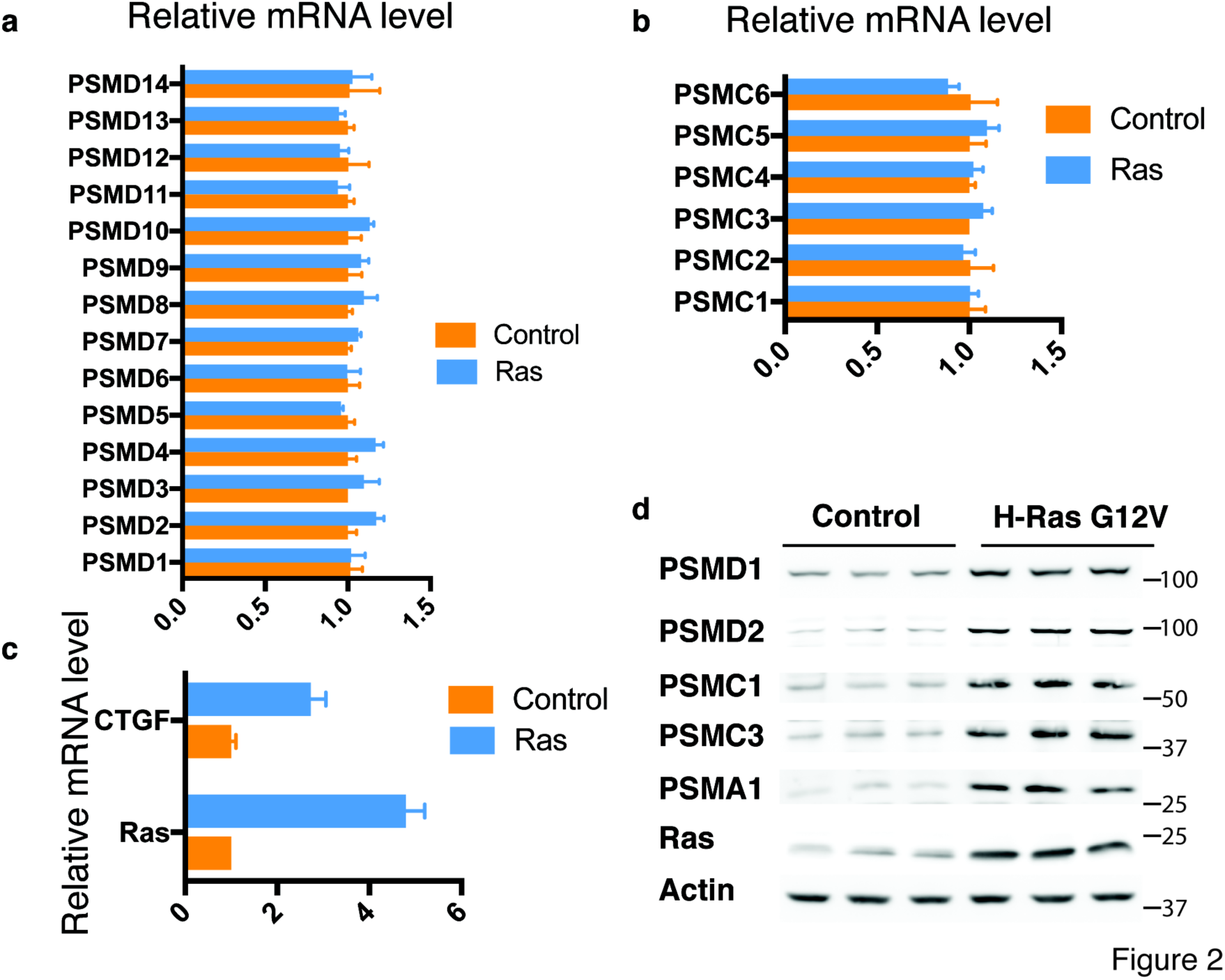
H-Ras G12V-transformed MCF10A cells accumulate proteasome subunits post-transcriptionally. MCF10A were transduced with retroviruses carrying pBabe H-Ras G12V gene. mRNA levels of all PSMD **(a)** and PSMC **(b)** proteasomal subunits, as well as of Ras and CTGF genes **(c)**, were measured by qPCR. Protein levels of selected proteasomal subunits were tested in triplicate by immunoblot **(d)**.

### H-Ras G12V transformed MCF10A are addicted to high 26S proteasome levels

Given the high 26S proteasome levels in H-Ras V12-transformed cell lines, we set out to explore the degree of dependency of the transformed state on high 26S proteasome levels. To do so, we established an isogenic model of H-Ras V12 transformation where specific reduction of 26S proteasomes is achieved by transient overexpression of an shRNA targeting the 19S subunit PSMD1. We first transduced MCF10A cells with a lentivector (LV) expressing a doxycycline inducible shRNA targeting the 19S subunit PSMD1 (Figure 3a). Cells were then transformed by over-expression of the oncogenic H-Ras G12V gene. As expected, a higher level of PSMD1 was observed in the Ras-transformed cells (Figure 3b). In the Ras-transformed cells the level of the PSMD1 (Rpn2) subunit was markedly reduced upon shRNA expression but remained higher than that found in naïve cells which do not express the shRNA. Similar results were obtained at the level of the 26S proteasome complex (Figure 3c). We will refer to this process as 26S depletion. The specific 26S depletion had a profound and selective effect on the viability of the Ras-transformed cells (Figures 3d,e). The H-Ras V12-transformed cells were significantly more sensitive to this depletion than the naïve cells. This is in spite the fact that the overall levels of 26S proteasomes are higher in H-Ras V12-transformed cells (Figure 3c). Thus, the increased levels of 26S proteasomes in the H-Ras V12-transformed cells, is crucial for their survival hinting toward their oncogenic addiction.

**Figure 3:**
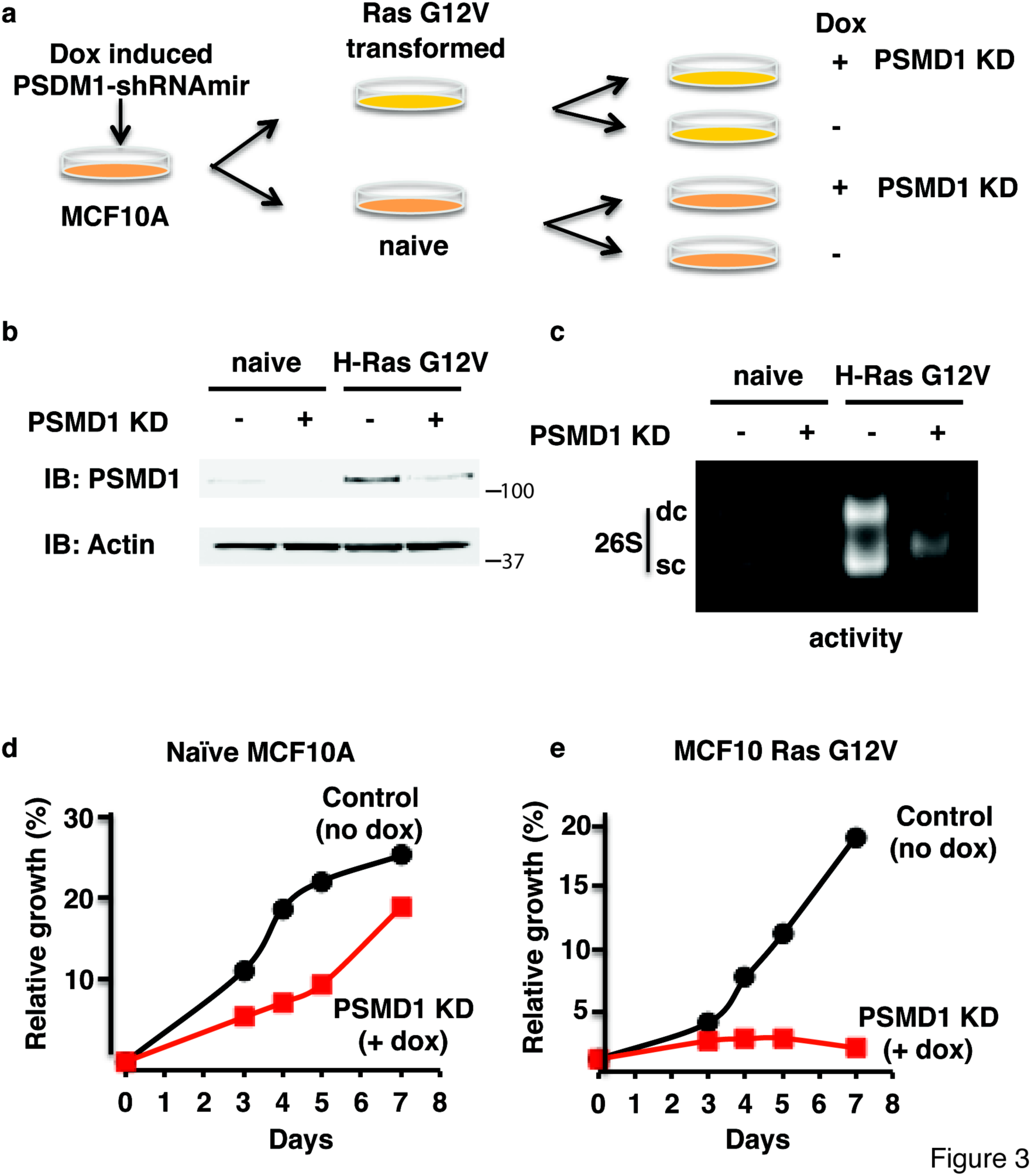
H-Ras G12V transformed MCF10A are addicted to high 26S proteasome levels. **(a)** MCF10A and MDA-MB-231 cells harboring a doxycycline-inducible PSMD1 shRNA were either doxycycline-treated to induce PSMD1 shRNA expression or left untreated. The levels of p53, p21 and PSMD1 were analyzed by immunoblot. **(b)** Growth of MDA-MB-231 cells expressing doxycycline-inducible PSMD1 shRNA was analyzed using the XTT assay. **(c)** Growth of MDA-MB-231 cells expressing doxycycline-inducible shRNA designed against the luciferase gene was analyzed as in b, ruling out non-specific doxycycline effects. **(d)** Visualization of the 26S-depleted MDA-MB-231 cells four days after PSMD1 shRNA induction. Expression of RFP as a marker for shRNA expression, and nuclear staining by DAPI are shown. Growth of MDA-MB-231 cells expressing doxycycline-inducible PSMD6 shRNA **(e)** and PSMD11 shRNA **(f)** was analyzed as in b. **(g)** Effect of synthetic siRNA designed against PSMD1, 6 or 11 or the luciferase gene on growth of MDA-MB-231 cells. Cells were transfected with the above siRNAs, and, after 24h were re-plated for the XTT assay. **(h)** Irreversible inhibition of cell proliferation and induction of cell killing in MDA-MB-231 cells by PSMD1 shRNA expression. Cells were induced to express PSMD1 shRNA for three days by doxycycline, and then washed and cultured in fresh medium without or with doxycycline for an additional 7 days. Cell proliferation was analyzed as in 4b.

### Addiction to high 26S proteasome levels in a triple negative breast cancer cell line

Having demonstrated that H-Ras V12-transformed cells are addicted to the 26S proteasome we next asked whether this is also the case with MDA-MB-231, a triple negative breast tumor cell line. Utilizing the same PSMD1 shRNA inducible strategy described above, a marked reduction at the level of the PSMD1 levels was obtained in these cells and the control MCF10A cells (Figure 4a). As expected, the level of p53 is high in MDA-MB-231 cells (41) but was not affected by 26S depletion. The level of p21 was increased in response to 26S depletion, which may result in cell cycle arrest. Since the MDA-MB-231 cells have a mutant p53, the obtained effect on cell viability is p53-independent. The reduction at the level of PSMD1 was sufficient to completely eliminate MDA-MB-231 cell growth (Figure 4b), whereas only a minor effect was observed on the MCF10A cells (Figure 3d). When shRNA designed against the Luciferase gene was induced by doxycycline cell growth was not affected (Figure 4c), ruling out non-specific doxycycline effects. Four days after 26S proteasome depletion only a few viable cells were observed exhibiting condensed nuclei (Figure 4d). Similar results were obtained with 26S depletion achieved by shRNAs targeting two other 19S complex subunits, namely PSMD6 (Figure 4e) and PSMD11 (Figure 4f) and with synthetic siRNA designed against each of these three subunits (Figure 4g). Both shRNA and siRNA designed against each of these three subunits were effective in knocking down the respective PSMD mRNA levels (Supplementary Figure 1a,b). Furthermore, the detrimental effect of 26S proteasome depletion in MDA-MB-231 cells was irreversible. PSMD1 shRNA expression for only three days was sufficient to induce irreversible inhibition of cell proliferation and induction of cell killing (Figure 4h). These data suggest that the TNBC are highly addicted to the 19S regulatory subunits required to establish high levels of the 26S proteasome.

**Figure 4:**
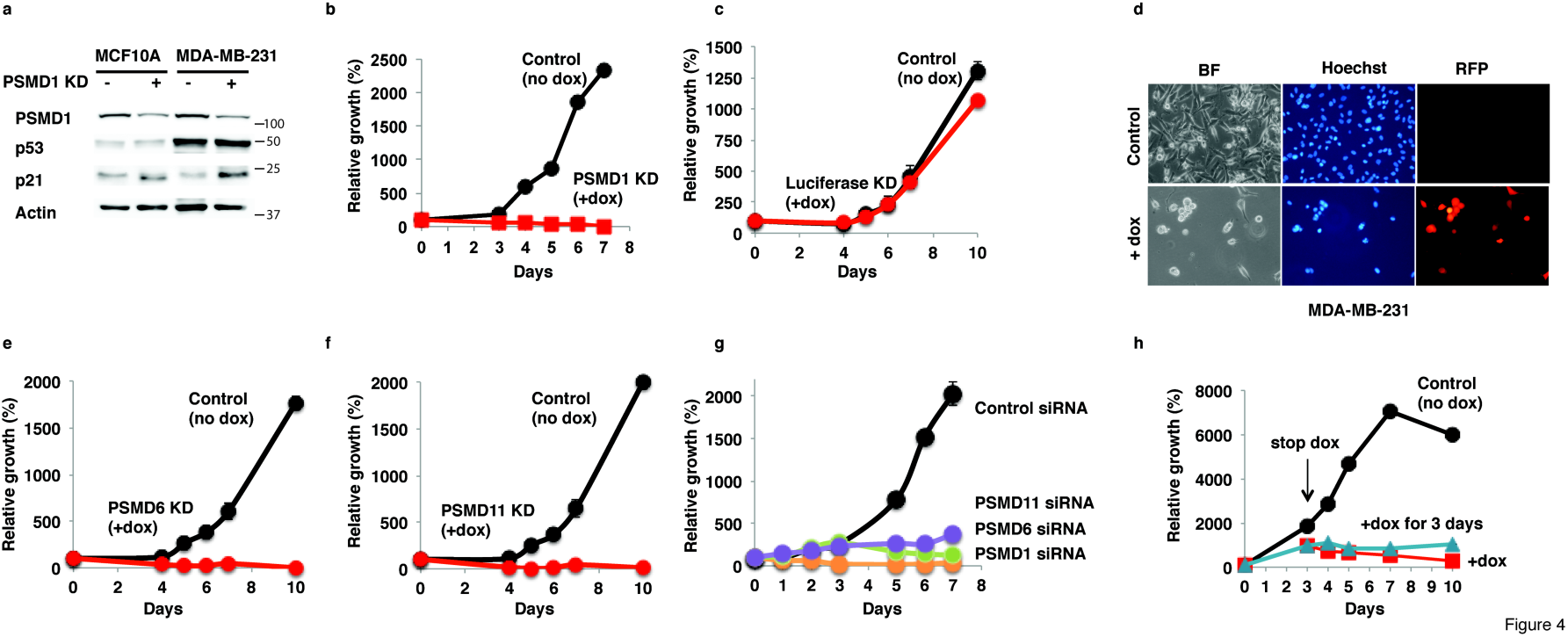
A triple negative breast cancer cell line is addicted to high 26S proteasome levels. **(a)** MCF10A and MDA-MB-231 cells harboring a doxycycline-inducible PSMD1 shRNA were either doxycycline-treated to induce PSMD1 shRNA expression or left untreated. The levels of p53, p21 and PSMD1 were analyzed by immunoblot. **(b)** Growth of MDA-MB-231 cells expressing doxycycline-inducible PSMD1 shRNA was analyzed using the XTT assay. **(c)** Growth of MDA-MB-231 cells expressing doxycycline-inducible shRNA designed against the luciferase gene was analyzed as in b, ruling out non-specific doxycycline effects. **(d)** Visualization of the 26S-depleted MDA-MB-231 cells four days after PSMD1 shRNA induction. Expression of RFP as a marker for shRNA expression, and nuclear staining by DAPI are shown. Growth of MDA-MB-231 cells expressing doxycycline-inducible PSMD6 shRNA **(e)** and PSMD11 shRNA **(f)** was analyzed as in b. **(g)** Effect of synthetic siRNA designed against PSMD1, 6 or 11 or the luciferase gene on growth of MDA-MB-231 cells. Cells were transfected with the above siRNAs, and, after 24h were re-plated for the XTT assay. **(h)** Irreversible inhibition of cell proliferation and induction of cell killing in MDA-MB-231 cells by PSMD1 shRNA expression. Cells were induced to express PSMD1 shRNA for three days by doxycycline, and then washed and cultured in fresh medium without or with doxycycline for an additional 7 days. Cell proliferation was analyzed as in 4b.

### Aggressive and drug resistant tumor cell lines are more susceptible to 26S depletion

We next asked how general is the 26S proteasome addiction across numerous cancer lines from distinct lineages. We utilized our experimental strategy of depletion of a 19S subunit in normal human foreskin fibroblast (HFF) and in multiple cancer cell lines. In all tested cells the PSMD1 subunit suppression was successful (Supplementary Figure 2). The viability of the normal HFF cells was not compromised under this condition (Supplementary Figures 3,4). In contrast, the viability of all tested transformed cells was either partially or completely compromised (Supplementary Figure 4). Heat map analysis revealed that many cell lines were extremely susceptible to this treatment (Figure 5a). We compared the immortalized MCF10A to its Ras-transformed counterpart, to demonstrate the power of our analysis in highlighting the effect of transformation.

**Figure 5:**
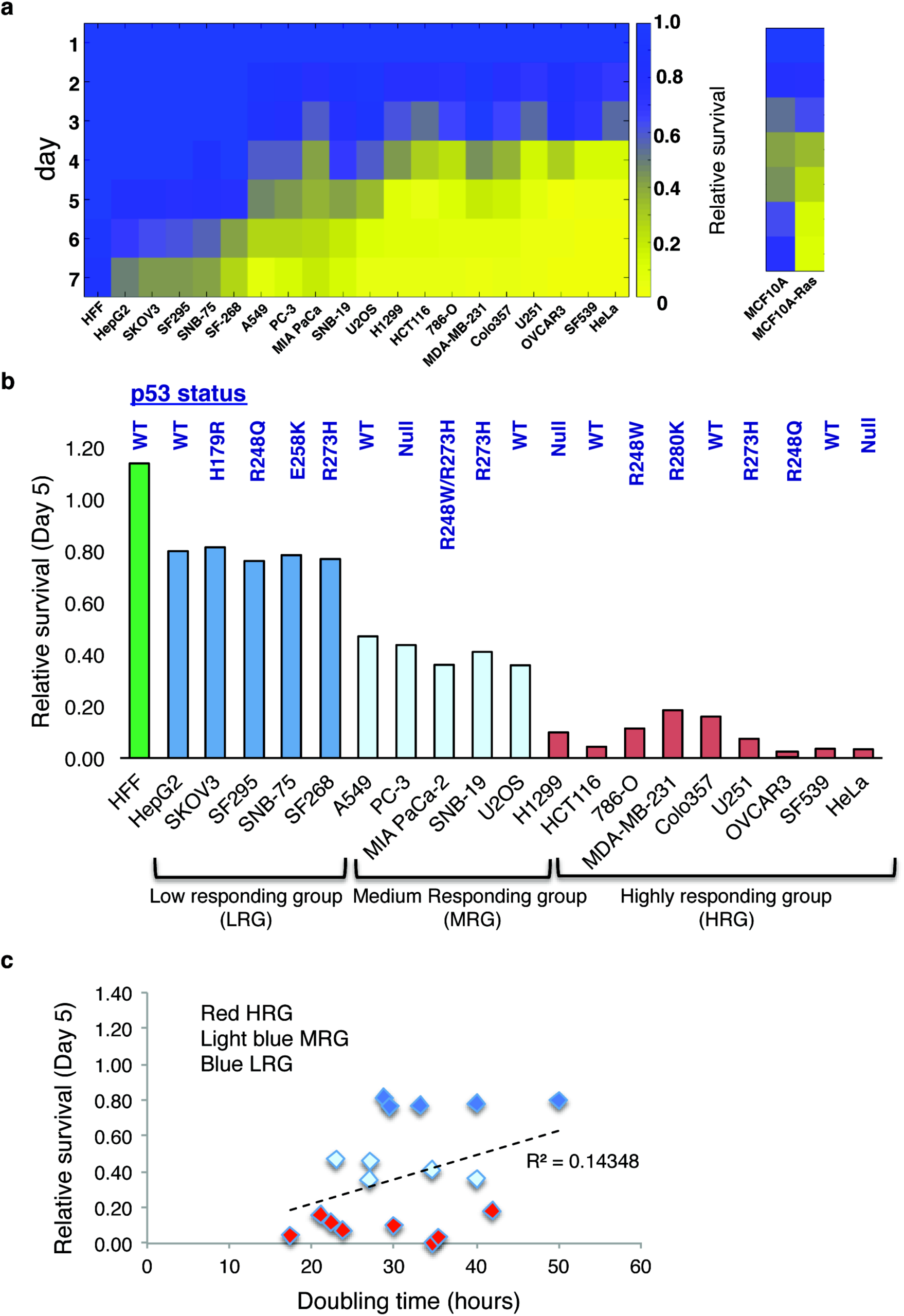
Aggressive and drug-resistant tumor cell lines are more susceptible to 26S depletion. A panel of various cell lines harboring doxycycline-inducible PSMD1 shRNA were treated with doxycycline to induce PSMD1 shRNA expression or left untreated. Alternatively, cells were transfected with siRNA against PSMD1 or luciferase gene. Cell proliferation was quantified as described in Figure 3d. **(a)** heat map analysis showing the relative survival of multiple cell lines in the course of 26S depletion (left). Heat map analysis showing relative survival of an isogenic cell line pair (MCF10A versus Ras-transformed counterpart) (right). Relative survival is calculated as a ratio of viability of PSMD1 shRNA-expressing cells to the viability of control cells measured at the same time point. **(b)** Relative survival of different cell lines measured at day 5 of induction. Status of the p53 gene is depicted for each cell line. **(c)** Relative survival at day 5 plotted against the doubling time of the tested cell lines.

Next, in order to compare between the cell lines, we plotted the response at day 5 and based on this analysis we divided the cell lines into four categories (Figure 5b). These include the normal non-responding HFF cells, the low responding, medium responding and the high responding cell lines. The high responding group includes U251, OVCAR3 and MDA-MB-231 that are all drug resistant and Colo357 and H1299 that were derived from metastatic sites (ATCC). Like MDA-MB-231, Colo321 and HCT116 were also vulnerable to 26S depletion achieved by PSMD6 and PSMD11 suppression (Supplementary Figure 5). Interestingly, there is no correlation between the status of the p53 gene and the level of viability in response to 26S depletion (Figure 5b). These data suggest that the more aggressive cancer lines are the more vulnerable they are to specific 26S proteasome depletion mediated by knocking down of the PSMD 19S particle subunits.

### Doubling time does not correlate with increased 26S depletion susceptibility

The selective killing of the tumor cells upon 26S depletion could be the result of the unique cell cycle doubling time of each of the lines. Using the reported doubling time (http://www.nexcelom.com/Applications/bright-field-analysis-of-nci-60-cancer-cell-lines.php) of the examined cell lines we plotted the survival ratio against the doubling time and found no correlation (Figure 5c). These data suggest that the mode of response of the tumor cell lines to 26S depletion is not the outcome of their rate of propagation.

### UPR activation in 26S depleted cells

To address the question of the mechanism of the decreased viability of the cell lines of the “high responding group”, we investigated the activation of the unfolded protein response (UPR). As expected, we found that 26S depletion increased the level of ubiquitinated proteins (Figures 6a-c). This accumulation is expected to induce UPR. ATF4 accumulation is a well-known hallmark of UPR activation, interestingly however, the level of ATF4 was not increased much in Colo357 and MDA-MB-231cells upon 26S depletion (Figures 6a and c). These cells also were rather poor in inducing sXBP-1 level (Figures 6d,e). In contrast, the level of phosphorylated eIF-α the upstream ATF4 regulator, was increased in all the three tested cell lines (Figures 6a-c). Also, the level of CHOP mRNA, the downstream target of UPR was increased in all the tested cell lines (Figures 6d-f). Thus, 26S depletion activates the UPR inducing the high expression of the apoptotic gene CHOP, which is expected to induce apoptosis.

**Figure 6:**
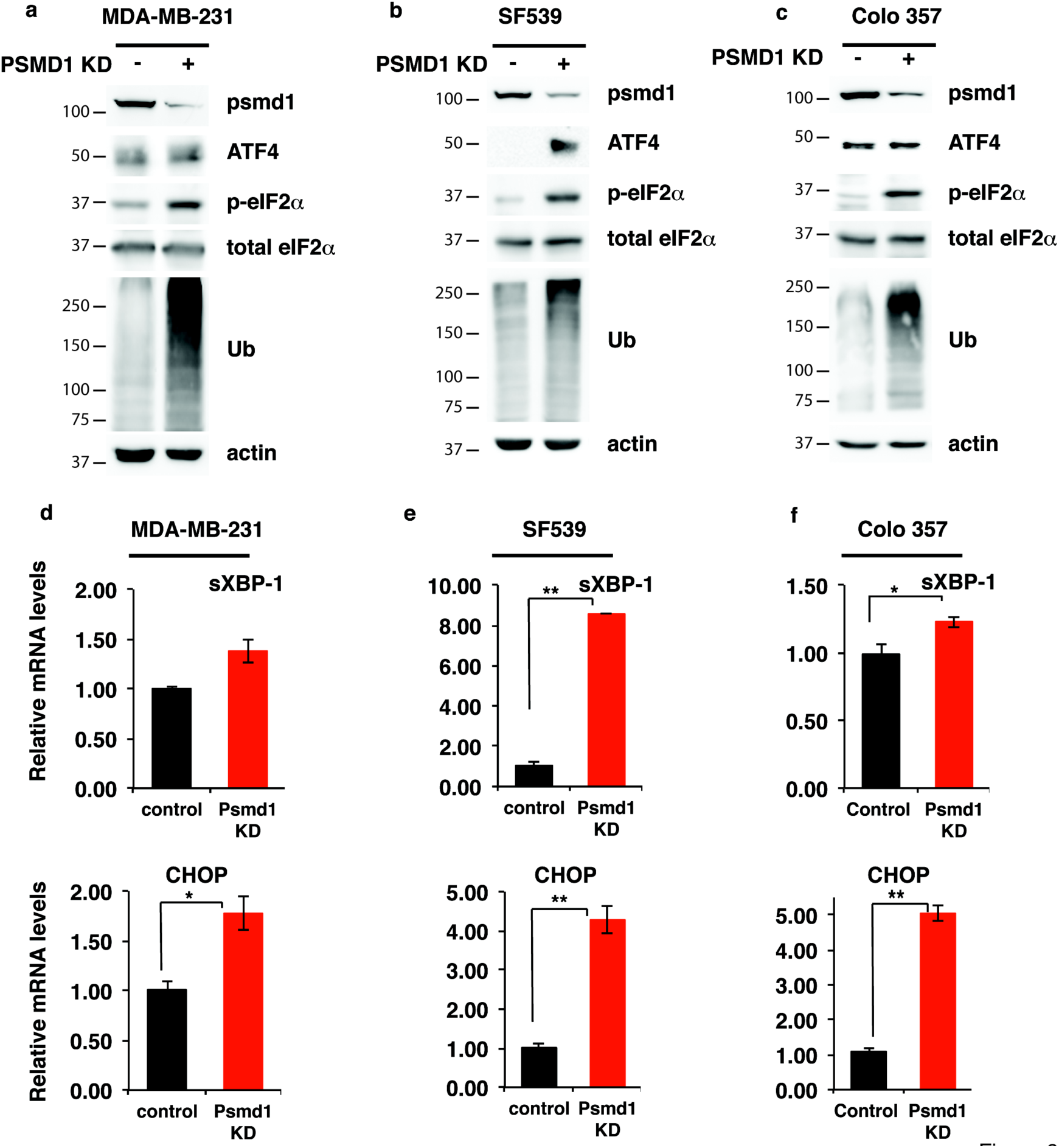
UPR activation in 26S depleted cells. Representative cell lines from the “highly responding group” MDA-MB-231 **(a, d)**, SF539 **(b, e)** and Colo357 **(c, f)** harboring doxycycline-inducible PSMD1 shRNA were either doxycycline treated to induce PSMD1 shRNA expression or left untreated. The levels of UPR markers ATF4 and p-eIFα were analyzed by immunoblot, and of CHOP and the spliced XBP-1 (sXBP-1) mRNA were measured by qPCR. * p<0.05, ** p<0.01.

### 26S depletion induces cytosolic condensation and nuclear distortion

To visualize the effect of the 26S depletion on cellular proteasome localization at single cell resolution we utilized CRISPR/Cas9 editing to tag the endogenous 20S proteasome complexes with YFP (Figure 7a). This did not affect the ability of 26S depletion to induce cell death (Figures 7b, c). Overall, the 26S proteasomes were localized to the cytoplasm and 26S depletion did not alter this cytoplasmic localization. However, 26S-depleted cells exhibited a rounded morphology with condensed cytoplasm and nuclear deformation and fragmentation (Figure 7d). This nuclear morphology is often observed in cells undergoing apoptosis (42). These data suggest that nuclear fragmentation is a likely mechanism of cell death in 26S-depleted cell lines.

**Figure 7:**
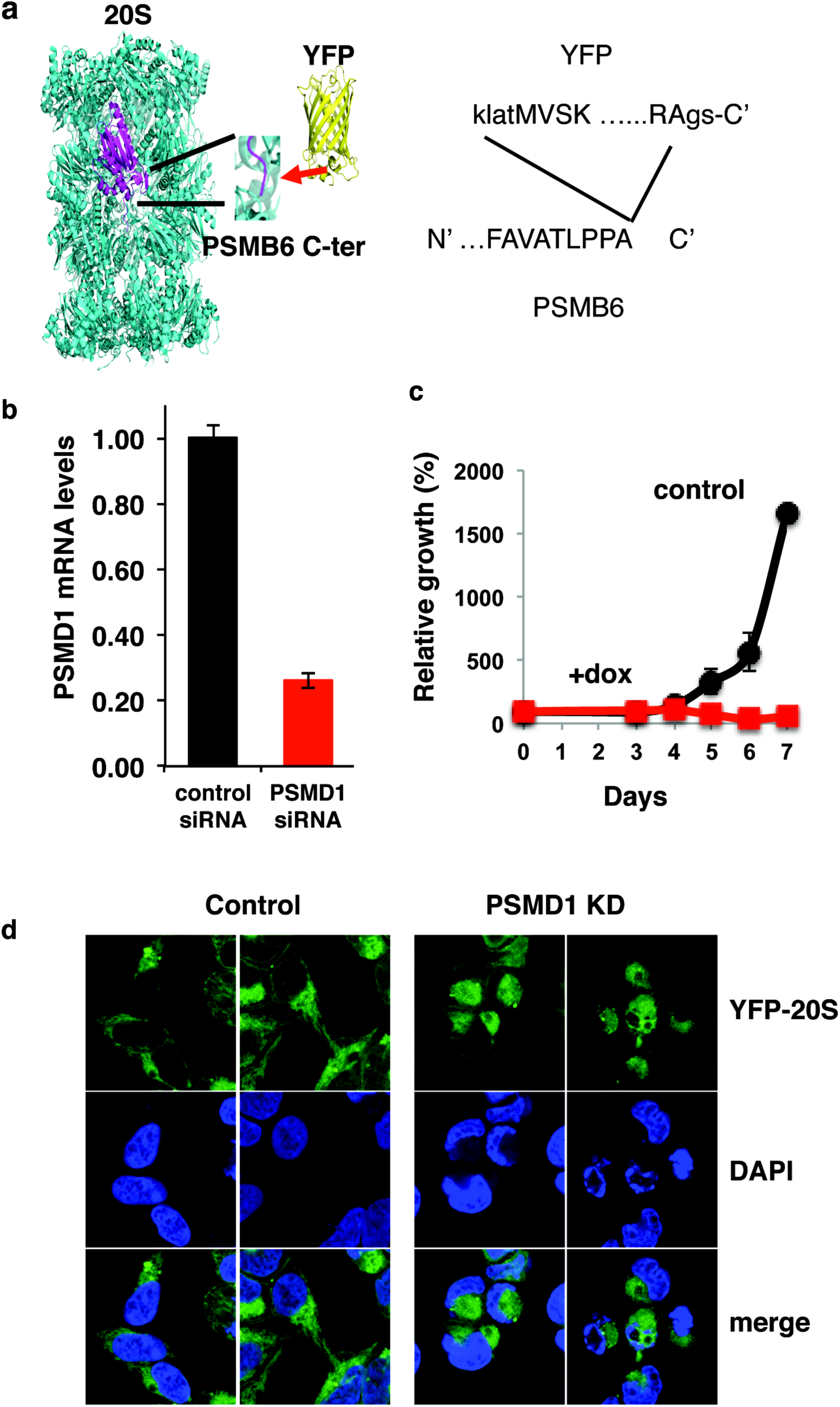
26S depletion induces cytosolic condensation and nuclear distortion. **(a)** Experimental strategy of CRISPR/Cas9 editing to tag the endogenous 20S subunit PSMB6 with YFP at its C-terminus (the N-and C-terminus of the YFP sequence are shown by capital letters. **(b-d)** HEK293 PSMB6-YFP cells were transfected with siRNA targeting PSMD1 or with control siRNA targeting luciferase. **(b)** PSMD1 levels in siRNA transfected PSMB6-YFP HEK293 cells were measured by qPCR. **(c)** growth of PSMB6-YFP HEK293 cells transfected with PSMD1 or control siRNA. **(d)** Cellular morphology of PSMB6-YFP 293 cells upon 26S depletion. Endogenous proteasomes are visualized by YFP, and nuclei are stained by DAPI.

### The role of Caspases in 26S depletion-mediated TNBC cell death

Cells undergoing apoptosis via nuclear fragmentation can be detected by measuring the subG1 fraction of the cell cycle. Unlike the naïve MCF10A cells, the Ras-transformed counterpart become enriched in subG1 fraction upon 26S depletion (Figures 8a, b). This is the case also with the MDA-MB-231 TNBC cell line (Figure 8c). Nuclear condensation and fragmentation is expected to induce Caspase 3 cleavage. Our data confirmed this prediction and showed that Caspase 3 is selectively activated in the TNBC cell line (Figure 8d). We next used QVD, the pan caspase inhibitor to test the role of caspases in cell death mediated by 26S depletion. As expected, QVD inhibited caspase cleavage/activation but unexpectedly it also diminished γH2AX activation (Figure 8e). Under this condition, the level of the accumulated polyubiquitinated proteins in the 26S depleted cells was not affected (Figure 8e). QVD treatment had a significant effect in preventing cell death following 26S depletion. However, inhibiting the caspase activity had only a partial rescue phenotype (Figure 8f). These data suggest that 26S depletion preferentially kills tumor cell lines partially via the induction of caspase-mediated cell death.

**Figure 8:**
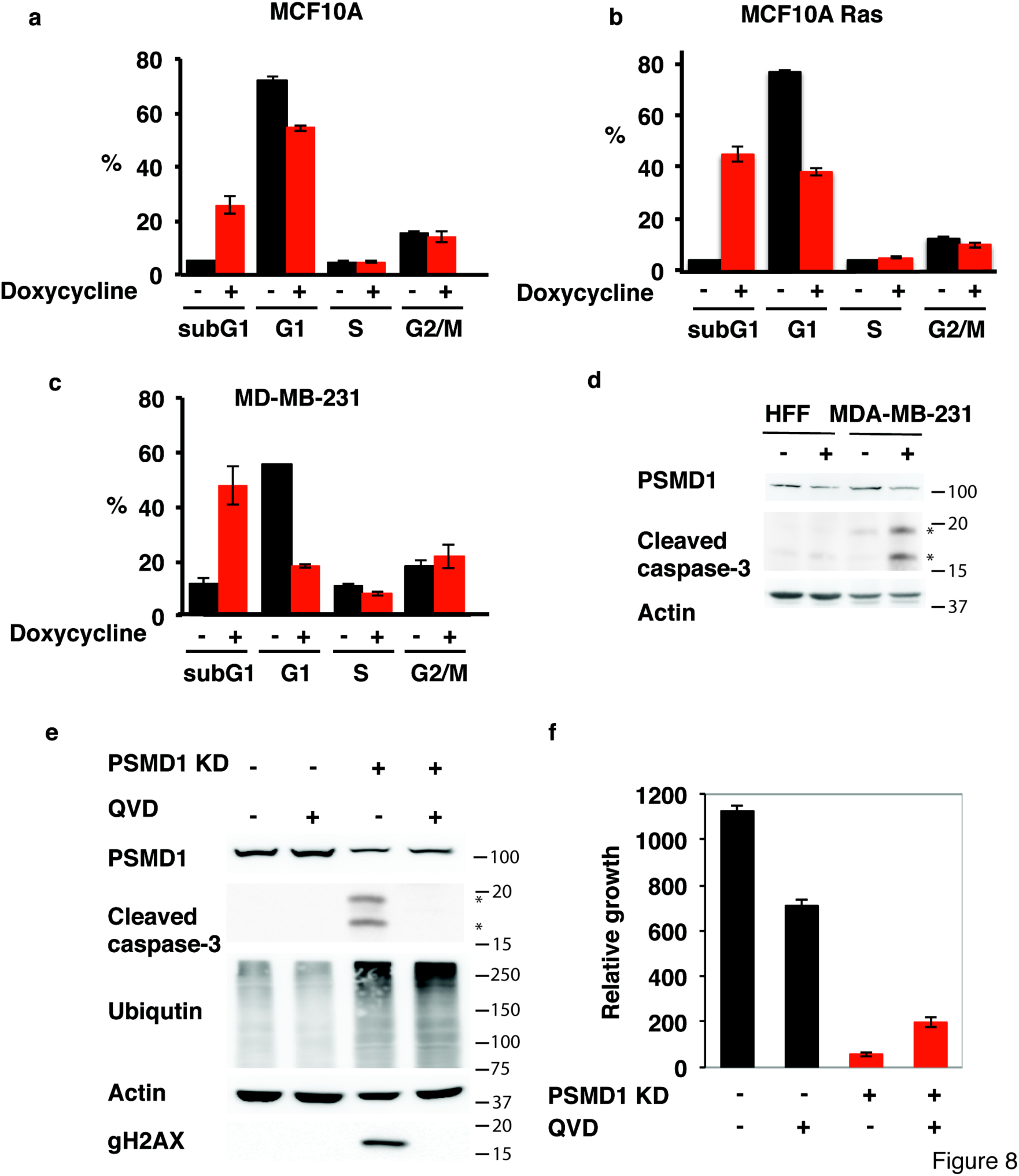
The role of caspases in 26S depletion-mediated TNBC cell death. Differential cell death response of **(a)** naïve MCF10A, **(b)** Ras-transformed MCF10A, and **(c)** cancer cell line MDA-MB-231 upon 26S depletion. Cells harboring doxycycline-inducible PSMD1 shRNA were either doxycycline-treated to induce PSMD1 shRNA expression or left untreated. Cell cycle was analyzed by FACS after 4 days, and cell death level was quantified by measuring subG1 fraction**. (d)** caspase-3 cleavage in MDA-MB-231 cells versus normal fibroblasts was analyzed by immunoblot after 4 days of PSMD1 shRNA induction. **(e)** role of caspases in cell death mediated by 26S depletion was examined by using pan-caspases inhibitor QVD. Cells were induced to express PSMD1 shRNA by doxycycline in the presence or absence of 25 µM QVD, and protein expression **(e)** and viability **(f)** were analysed after 4 days.

## Discussion

We report here that transformation by H-Ras G12V of the mouse NIH3T3 and human MCF10A immortalized cell lines is associated with a marked increase in the level of the 26S proteasome complexes. During the process of transformation there is increased dependency on proteasome function as part of global increased burden on the protein homeostasis machinery. Genetic screen analysis in several models including Ras transformation and triple negative breast cancer cells revealed a strong dependency on the proteasome function (38–40). Our analysis revealed that Ras-transformed cells cannot survive with the amount of the 26S proteasome that the naïve MCF10A cells normally have. Thus, the high 26S proteasome level is in fact critical for the survival of the transformed cells.

Reducing the expression of one of the key 19S subunits (with shRNA or siRNA) results in reduction of 26S proteasome levels and activity and effective inhibition of the ubiquitin-dependent degradation process, while leaving the 20S catalytic particle intact and active (7). Utilizing this experimental strategy, we characterized the dependency of cancer cells on specific 26S proteasome function. This is not only the case with the Ras-transformed cells but holds across numerous cancer cell lines. The 19S subunits are essential (43), therefore, complete suppression would result in cell death. However, our findings reveal differential effects of partial 19S subunit suppression. Under normal non-oncogenic conditions cells can tolerate moderate inhibition of the 26S proteasome. Upon oncogenic transformation, despite having more 26S proteasomes cells become increasingly more sensitive to suppression of the 26S proteasome. This is in particular the case with the drug-resistant and more aggressive tumor cell lines. Thus, oncogenic transformation entails addiction to high 26S levels and function.

Targeting the proteasomes for cancer therapy proved to be effective mostly in multiple myeloma. Targeting the functional subunits of the 19S regulatory particle could serve as an attractive alternative approach with minor effects on the normal cells. Along this line, previous attempts were made to inhibit the substrate recognition and deubiquitination activity of the 19S particle components. RPN13 is one of two major ubiquitin receptors within the 19S regulatory particle. The small molecule RA190 covalently binds to RPN13 and inhibits its function and is active against cervical and ovarian cancer (44). The 19S particle further contains RPN11, and two cysteine proteases USP14 and UCHL5 (UCH37) that have de-ubiquitinating (DUB) activity. DUB inhibitors were shown to inhibit tumor progression in certain solid tumor models (45, 46). Our finding that the transformed cells accumulate high amounts of the 26S proteasome and that depletion of a single 19S particle subunit reduces the 26S proteasome level, set the stage for a new strategy of cancer therapy. This strategy does not deal with the 19S particle associated activities but rather depletes the 26S complex. The observation that the normal cells are refractory to this process suggest that a very low amount of the 26S proteasome is required to survive.

A critical question is why the tumor cells are addicted to high 26S proteasome level, a behavior that is positively correlated with aggressiveness of the tested cell lines. A likely speculation is that it is due to increased protein homeostasis burden, proteotoxic stress, that is associated with cancer transformation (39, 47, 48). A number of mechanisms have been proposed in explaining the cytotoxic effects of proteasome inhibitors. These include NF B inhibition (49), expression of Myc (50), or accumulation of the proapoptotic proteins (Bim, Bik and Noxa) (50, 51). However, it is rather likely that these processes are the result of the activation of the unfolded protein response (UPR) (52). In our tested cell lines UPR was activated upon 26S proteasome depletion but there is no strong correlation between the level of UPR activation and cell viability. The treated cells show accumulation of subG1 fragmented cells and nuclear deformation which are the hallmark of Caspase activation and DNA fragmentation. However, the pan Caspase inhibitor only partially rescued cell death. Furthermore, antioxidants failed to rescue the cells as well (not shown). Therefore, the robust mechanism regulating cancer cell death under 26S proteasome depletion still needs to be better characterized.

The 26S proteasome is the end point of the ubiquitin proteasome pathway that is chiefly required for cell cycle progression. It was therefore postulated, that the proliferation of cancer cells make them susceptible to proteasome inhibition. Our analysis revealed a lack of correlation between the 26S proteasome dependency and rate of cell proliferation. Moreover, in an isogenic model of Ras-transformed cells, where no significant increase in rate of proliferation was detected, a strong increase in 26S proteasome levels and dependency was observed suggesting other mechanism(s) may be in play. This brings forward the possibility that cancer cells may exhibit altered dependencies on the 26S proteasome function that is not associated with cell cycle progression.

A model that can be suggested to explain our findings is a 26S proteasome buffer model. This model shows strong resemblance to the buffer model described for the essential chaperone Hsp90. The levels of Hsp90 in the cell are constitutively higher than required to fulfill its normal function in the cell creating a reservoir-buffer (53). However, once protein homeostasis is perturbed by environmental or genetic insults the need and dependency on Hsp90 is increased due to proteotoxic stress, diminishing the access buffer of Hsp90 resulting in phenotypic outcome (54–56). In an analogous manner, our results support the model that there is a reservoir of 26S proteasomes in the cell. Under normal conditions the reduction of 26S proteasome levels is tolerated. However, upon oncogenic transformation the 26S proteasome buffer diminishes resulting in an increased dependency on elevated levels of 26S proteasome. Sampling different cancer cells we show here that this dependency varies between the different cells. Moreover, in some cancer cells and tumors there is naturally occurring (epi)genetic reduction in expression of one of the 19S subunits suggesting that some cancers can tolerate a mild reduction in the 26S proteasome buffer (26, 57). Thus, the 26S proteasome buffer model suggests that in the context of oncogenic transformation or in the case of more aggressive cancer cell lines where the protein homeostasis burden is expected to be increased, the dependency on 26S proteasome would be augmented as the 26S buffer would be depleted increasing the vulnerability to targeted 26S suppression.

## Materials and methods

### Cells

MCF10A cells were cultured in DMEM: F12 medium with 5% horse serum (Gibco), 2 mM glutamine, 20 ng/ml epidermal growth factor, 10 µg/ml insulin, 0.5 mg/ml hydrocortisone, 100 ng/ml cholera toxin,100 units/ml penicillin and 100 µg/ml streptomycin. OVCAR3, SKOV-3, MDA-MB-231, H1299, A549, 786-O, U251, SF268, SF295, SF539, SNB-75, SNB-19, Colo357 cells were cultured in RPMI-1640 medium (Gibco) with 10% FCS 2mM glutamine and the antibiotics as above. HEK 293T, NIH3T3, HCT116, U2OS, HepG2, PC-3, HEK293, and MiaPaCa-2 were cultured in DMEM and 8% FCS with antibiotics as mentioned above. Human foreskin fibroblasts (HFF) were cultured in DMEM with 10%FCS, 2mM glutamine and the antibiotics as above. All cells were maintained at 37°C in a humidified incubator with 5.6% CO_2_.

### Plasmids and viral production

A lentiviral Tet-inducible TRIPZ vector with shRNAmir against 26S proteasome subunit PSMD1 was purchased from Open Biosystems (Thermo Scientific) and used to down regulate 26S levels. To down regulate PSMD6 and PSMD11 26S proteasome subunits, shRNA targeting the sequence 5′-CAGGAACTGTCCAGGTTTATT -3′ (for PSMD6) or 5′-GGACATGCAGTCGGGTATTAT -3′ (for PSMD11) were cloned into the Tet-pLKO-puro inducible vector (Addgene plasmid #21915). Transducing lentiviral particles were produced in HEK293T cells according to the manufacturer's protocol. For H-Ras transformation of NIH3T3 cells, production of retrovirus particles was performed in HEK 293T cells with either pBabe empty or pBabe H-Ras G12V and psi helper plasmid. For H-Ras transformation of MCF10A cells, production of retrovirus particles was done in the Phoenix packaging cell line (kindly provided by Dr. Gary Nolan, Stanford University, Stanford, CA) with either pBabe H-Ras V12 vector or an empty pBabe vector.

### Induction of PSMD1 knockdown and reduction in the 26S proteasomal complexes

Cells of interest were infected with the lentivirus particles (infection conditions vary between cell lines). Selection with puromycin was employed for a week (the concentration has to be determined based on the cell type). To induce shRNA expression to knockdown PSMD1 the cells were treated with 1 µg/ml doxycycline. The efficiency of the reduction in the 26S proteasomal complex is analyzed by subjecting the cellular extract to nondenaturing PAGE as previously described (58). This enables visualization of the reduction in the 26S proteasomal complex and not only the subunit expression on the protein level (as examined by the SDS-PAGE).

### Flow cytometry

Cells were seeded at a density of 5×10^5^ per 9 cm dish and PSMD1 knockdown was induced by the addition of doxycycline (1 µg/ml) in the presence of puromycin (2 µg/ml). After 96 hours floating and attached cells were collected and combined together, washed twice with PBS and fixed in 70% ethanol. The cells were further washed with PBS and resuspended in 50 µg/ml RNase A and 25 µg/ml propidium iodide in PBS. In each assay, triplicates of 30,000-50,000 cells were collected by the BD LSRII flow cytometer and analyzed with the BD FACSDiva software (BD Biosciences).

### Cell proliferation

Cells (2×10^3^) were seeded in a well of 96-well plate in the presence of puromycin (2 µg/ml) with or without doxycycline (1 µg/ml). Cell proliferation was analyzed using the XTT assay (Biological Industries) and spectrophotometrically quantified. The results obtained by the XTT assay were confirmed by direct counting of cells over the period of PSMD1 KD induction.

In the recovery experiment, MDA-MB-231 cells (2 × 10^3^) were seeded in a well of 96-well plate in the presence of puromycin (2 µg /ml) and doxycycline (1 µg /ml). After 72 h, the cells were washed twice with the culture medium and supplemented with fresh medium with puromycin but without doxycycline for an additional seven days. The ability of cells to recover from PSMD1 KD was compared to growth characteristics of cells that continued to be supplemented with 1 µg /ml doxycycline after the washing step or were not induced at all.

### Anchorage-independent growth

Cells (3×10^4^) were added to 0.5 ml of growth medium with 0.3% low-melting point agar (BioRad) and layered onto 0.5 ml of 0.5% agar beds in twelve-well plates in the presence of puromycin (2 µg/ml) with or without doxycycline (1 µg/ml). Colonies were photographed after one week.

### Colony-formation assay

Cells were seeded at a density of 30 cells/cm^2^ and cultured in the presence of puromycin (2 µg/ml) with or without doxycycline (1 µg/ml) for 14 days. Colonies were fixed with 70% isopropanol for 10 min followed by staining with 0.1% crystal violet.

### Protein extraction and immunoblot analysis

Cells were collected and lysed in RIPA buffer (50 mM Tris-HCl pH 8, 150 mM NaCl, 1% Nonidet P-40 (v/v), 0.5% deoxycholate (v/v), 0.1% SDS (v/v), 1mM DTT and Sigma protease inhibitor mixture (P8340) as previously described (59), the extract was subjected to ultracentrifugation (13,000g, 15min) and the supernatant was used as the protein extract. For detection of H2AX phosphorylation, whole cell lysates were prepared by incubation in RIPA buffer at 4°C for 20 min, followed by sonication. Protein concentrations were determined by Bradford assay (Bio-Rad) and samples were mixed with Laemmli sample buffer [final concentration 2% SDS, 10% glycerol, 5% 2-mercaptoethanol and 0.0625 M Tris-HCl pH6.8], boiled for 5 minutes and loaded on a polyacrylamide-SDS gel. Following electrophoresis, proteins were transferred to cellulose nitrate 0.45 µm membranes (Schleicher & Schuell). The primary antibodies used are cited in Sup. Table 1. Secondary antibodies were HRP-linked goat anti-mouse, goat anti-rabbit and donkey anti-goat antibodies (Jackson ImmunoResearch). Signals were developed using the Ez-ECL kit (Biological Industries) and detected by ImageQuant LAS 4000 (GE).

### Nondenaturing PAGE

Proteasomal samples were loaded on a non-denaturing 4% polyacrylamide gel using the protocol previously described (58, 60). Gels were either overlaid with Suc-LLVY-AMC (50µM) for assessment of proteasomal activity by ImageQuant LAS 4000 (GE) or transferred to nitrocellulose membranes where immunoblotting with anti PSMA4 antibody was conducted.

### RNA interference and microscopic studies

PSMB6-YFP HEK293 cells were transfected with 100 nM of RNAi targeting PSMD1 (directed against sequence 5’-CTCATATTGGGAATGCTTA-3’) or control RNAi targeting luciferase, by using Dharmafect 4 reagent of Dharmacon. On the next day, the transfected cells were re-plated on glass coverslips precoated with poly-lysine. Forty eight hours following transfection cells were fixed in 4% paraformaldehyde for 30 min and mounted in Aqua-PolyMount (Polysciences). Microscopic images were acquired using a Zeiss LSM 710 confocal scanning system, using ×63 NA (numerical aperture) 1.4 objectives, and processed using Adobe Photoshop CS6

### RNA extraction and analysis

Total RNA was extracted using the TRI Reagent (MRC). First strand synthesis was performed using iScript cDNA synthesis kit (Quanta). qRT-PCR was performed using the LightCycler480 (Roche), with PerfeCta®SYBR Green FastMix mix (Quanta). All qPCRs were normalized to TBP1 mRNA levels.

### Labeling of endogenous PSMB6 by YFP

For CRISPR/Cas9-mediated modification of endogenous proteasome subunit PSMB6, HEK293 cells were transfected with pX330-U6-Chimeric_BB-CBh-hSpCas9 (gift from Feng Zhang, Addgene plasmid # 42230) encoding sgRNA targeting the stop codon of PSMB6 (TAGAATCCCAGGATTCAGGC), and a donor plasmid with 1 kb homology arms flanking an SYFP insert. The resulting construct has SYFP fused to the C-terminus of PSMB6, with a 4 amino acid linker. The SYFP sequence was subcloned from pSYFP-C1 (gift from Dorus Gadella, Addgene plasmid #22878) using PCR.

## Acknowledgements

We thank Gary Nolan for the Phoenix packaging cell line. This research was supported by the Rising Tide Foundation and the Israel Science Foundation (grant No. 1591/15).

---

**Supplementary Figure 1.**
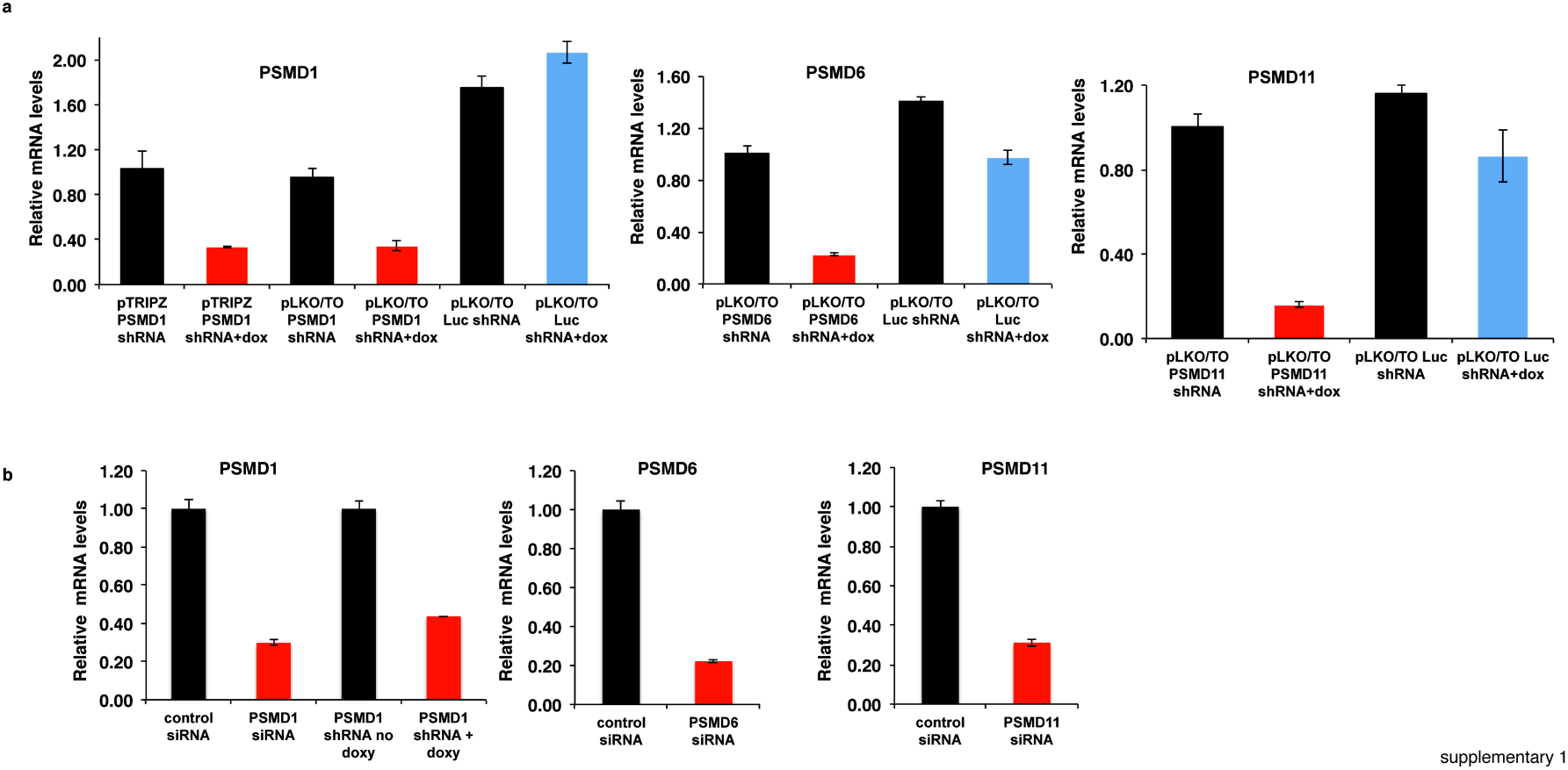
Quantification of shRNA-siRNA-mediated knockdown of 19S subunits PSMD1, PSMD6, and PSMD11. mRNA levels of PSMD1, PSMD6 and PSMD11 during 26S depletion were quantified by qPCR. **(a)** MDA-MB-231 cells harboring doxycycline-inducible PSMD1, 6, 11 shRNA or control luciferase-targeting shRNA were induced with doxycycline to express the respective shRNA for 3 days. **(b)** MDA-MB-231 cells were transfected with siRNA targeting PSMD 1, 6 and 11 or control luciferase-targeting siRNA. mRNA was analysed after three days by qPCR.

**Supplementary Figure 2.**
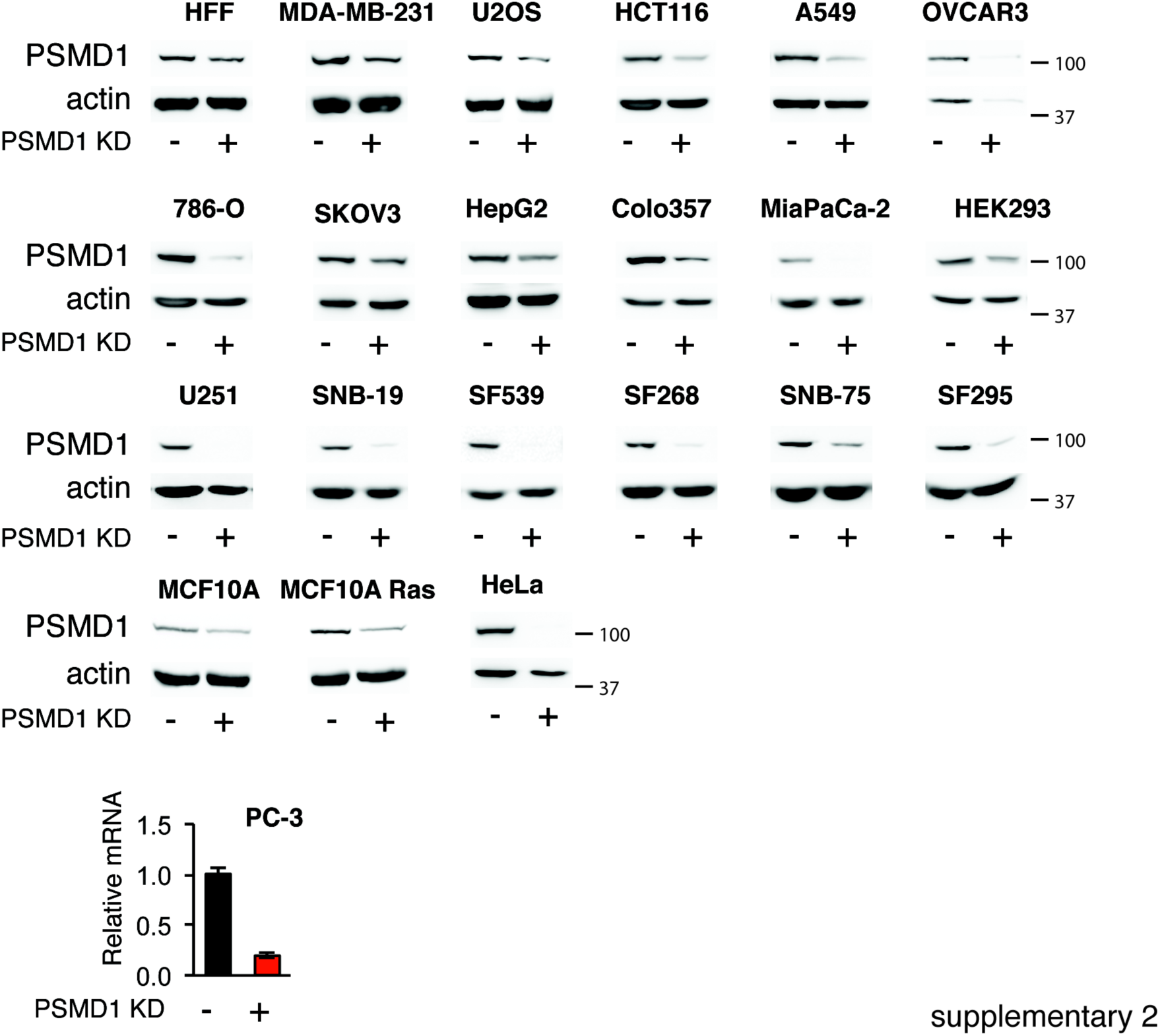
Reduction in PSMD1 levels during 26S depletion in various cell lines used in the study. Protein levels were analyzed by immunoblot, and mRNA levels were measured by qPCR after 3-4 days of 26S depletion.

**Supplementary Figure 3.**
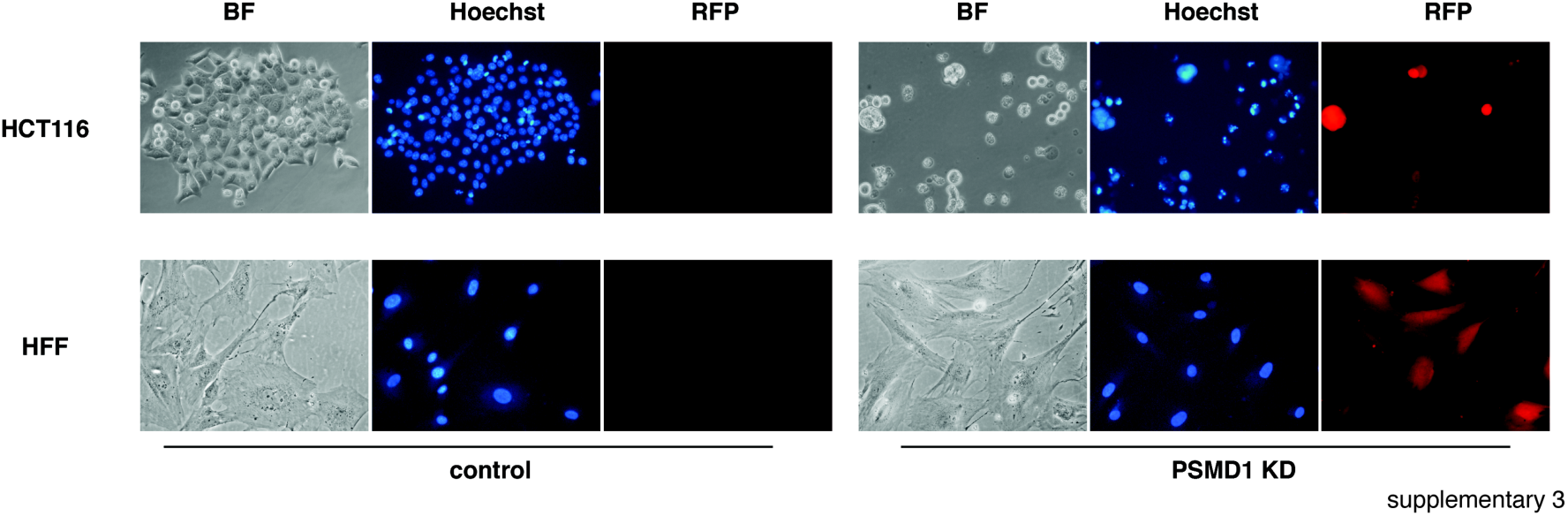
Nuclei deformations are observable in cancer cells (HCT116) but not in normal cells (HFF) with PSMD1 KD. Colon carcinoma (HCT116) and normal fibroblasts (HFF) were transduced using lentiviral PSMD1 shRNA expression vector. Cells were induced to express PSMD1 shRNA by doxycycline (1µg/ml). After 4 days of induction, nuclei were stained with Hoechst 33342 and images were taken using an inverted fluorescent microscope. Expression of RFP as a marker for shRNA induction and nuclear staining by DAPI are shown.

**Supplementary Figure 4.**
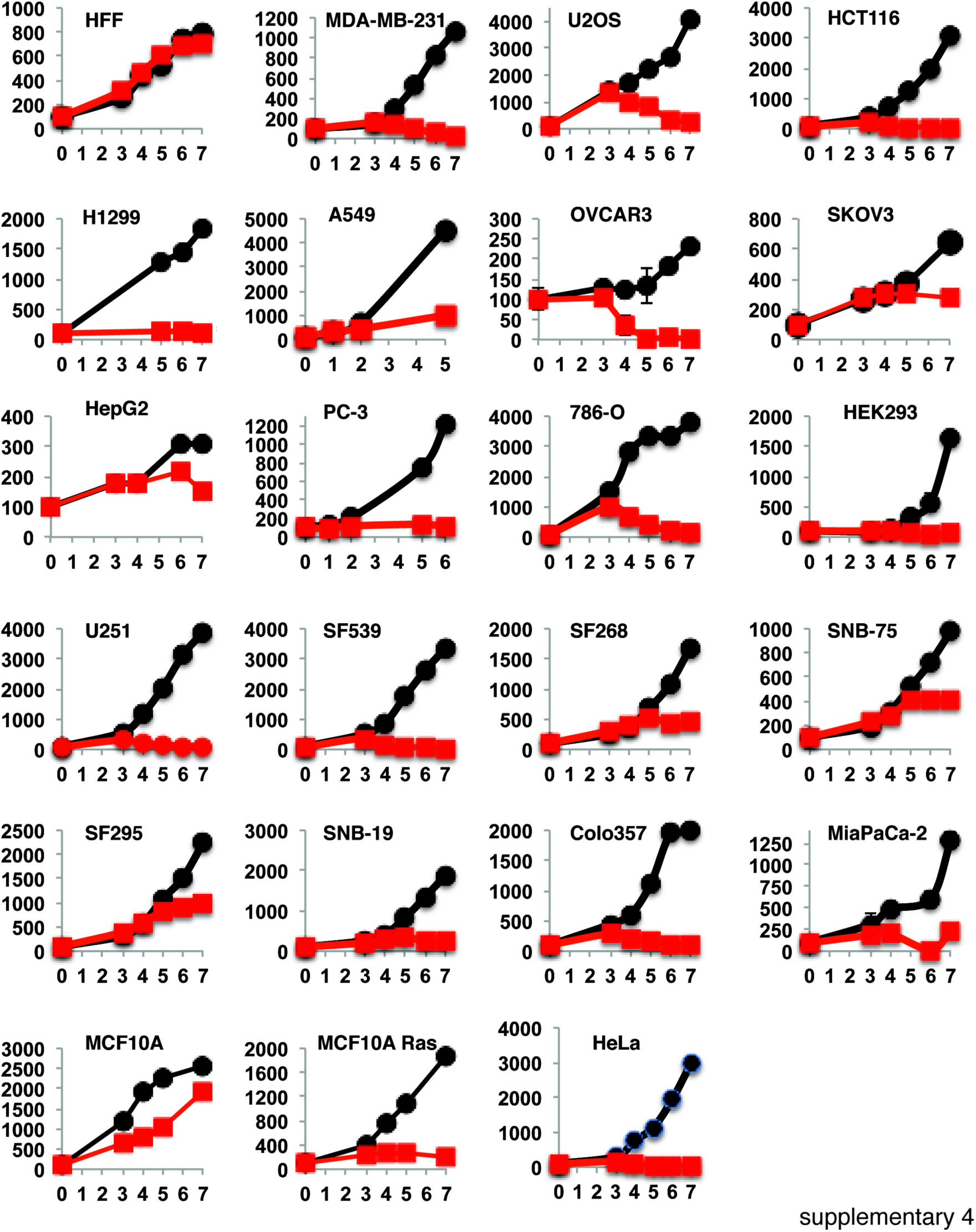
Relative cell growth of cell lines with and without PSMD1 knockdown. Growth of cell lines used in the study was measured as described in Figure 3d. Red, with, and black, without doxycycline induction of shRNA.

**Supplementary Figure 5.**
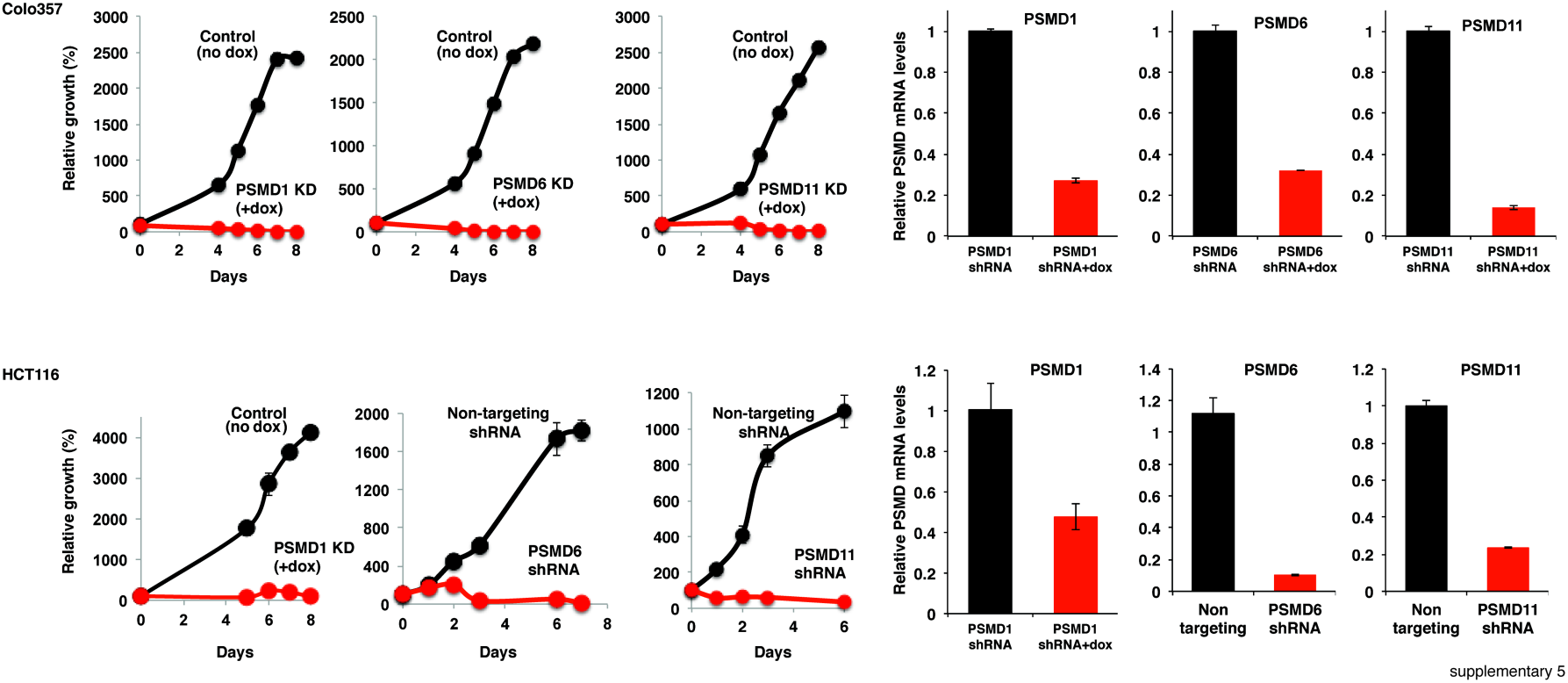
Vulnerability of Colo321 and HCT116 upon suppression of either PSMD1, PSMD6 or PSMD11 shRNA. Colo321 and HCT116 were transduced using lentiviral vectors for doxycycline inducible or constitutive expression of PSMD1, PSMD6, PSMD11 or control non-targeting shRNA. When appropriate, cells were induced to express shRNA by doxycycline (1µg/ml). Cell growth was analyzed by XTT, and the extent of knock down was tested by qPCR.

**Supplementary Table I:** Antibodies used

| Primary antibody | Source | Organism | Cat # |
| --- | --- | --- | --- |
| PSMD1 | Sigma | rabbit | S1324 |
| PSMA1 | Prof. Chaim Kahana, Weizmann Institute of Science | rabbit | - |
| PSMA4 | Prof. Chaim Kahana, Weizmann Institute of Science | rabbit | - |
| PSMD2 | Thermo Scientific | rabbit | PA1-964 |
| PSMC1 | Thermo Scientific | rabbit | PA1-965 |
| PSMC3 | Abnova | mouse | H00005702-B01P |
| Fibronectin | Sigma | mouse | F0916 |
| Myc | Weizmann Institute Ab Core Facility | mouse | - |
| c-Jun | Santa Cruz | rabbit |  |
| NQO1 | Santa Cruz | goat | C19, R20 |
| Hsc-70 | Prof. Irith Ginzburg, Weizmann Institute of Science | rabbit | - |
| actin | Sigma | mouse | A4700 |
| Beta-tubulin | Covance | mouse | MMS-435P |
| RNR-R2 | Santa Cruz | goat | sc-10844 |
| p53 | Prof. Moshe Oren,<br>Weizmann Institute of<br>Science | mouse | clone 1801 |
| p21 | Santa Cruz | Rabbit,<br>mouse | sc-397, sc-6246 |
| ATF4 | Cell Signaling | rabbit | #11815 |
| eIF2α | Santa Cruz | rabbit | sc-11386 |
| pSer52 eIF2α | Thermo Scientific | rabbit | 44-728G |
| ubiquitin | Santa Cruz | mouse | sc-8017 |
| cleaved caspase-3 | Cell Signaling | rabbit | #9664 |
| pSer139 H2A.X | Upstate Biotechnology | mouse | 05-636 |
| pan Ras | Santa Cruz | mouse | sc-166691 |

## References

1. Hershko A, Ciechanover A. The ubiquitin system. Annu Rev Biochem. 1998;67:425–79. Epub 1998/10/06.

2. Collins GA, Goldberg AL. The Logic of the 26S Proteasome. Cell. 2017;169(5):792–806.

3. Zwickl P, Voges D, Baumeister W. The proteasome: a macromolecular assembly designed for controlled proteolysis. Philos Trans R Soc Lond B Biol Sci. 1999;354(1389):1501–11.

4. Varshavsky A. The ubiquitin system. Trends Biochem Sci. 1997;22(10):383–7. Epub 1997/11/14.

5. Finley D, Chen X, Walters KJ. Gates, Channels, and Switches: Elements of the Proteasome Machine. Trends Biochem Sci. 2016;41(1):77–93.

6. Hochstrasser M. Ubiquitin-dependent protein degradation. Annu Rev Genet. 1996;30:405–39. Epub 1996/01/01.

7. Tsvetkov P, Reuven N, Prives C, Shaul Y. The susceptibility of the p53 unstructured N-terminus to 20S proteasomal degradation programs stress response. J Biol Chem. 2009.

8. Adler J, Reuven N, Kahana C, Shaul Y. c-Fos proteasomal degradation is activated by a default mechanism, and its regulation by NAD(P)H:quinone oxidoreductase 1 determines c-Fos serum response kinetics. Mol Cell Biol. 2010;30(15):3767–78.

9. Wiggins CM, Tsvetkov P, Johnson M, Joyce CL, Lamb CA, Bryant NJ, et al. BIM(EL), an intrinsically disordered protein, is degraded by 20S proteasomes in the absence of poly-ubiquitylation. J Cell Sci. 2011;124(Pt 6):969–77.

10. Shaul Y, Tsvetkov P, Reuven N. IDPs and Protein Degradation in the Cell: John Wiley & Sons, Inc.; 2010. 1–36 p.

11. Ben-Nissan G, Sharon M. Regulating the 20S proteasome ubiquitin independent degradation pathway. Biomolecules. 2014;4(3):862–84.

12. Asher G, Reuven N, Shaul Y. 20S proteasomes and protein degradation "by default". Bioessays. 2006;28(8):844–9.

13. Tsvetkov P, Reuven N, Shaul Y. The nanny model for IDPs. Nature chemical biology. 2009;5(11):778–81.

14. Hoeller D, Dikic I. Targeting the ubiquitin system in cancer therapy. Nature. 2009;458(7237):438–44. Epub 2009/03/28.

15. Bedford L, Lowe J, Dick LR, Mayer RJ, Brownell JE. Ubiquitin-like protein conjugation and the ubiquitin-proteasome system as drug targets. Nat Rev Drug Discov. 2011;10(1):29–46.

16. Navon A, Ciechanover A. The 26 S proteasome: from basic mechanisms to drug targeting. J Biol Chem. 2009;284(49):33713–8. Epub 2009/10/09.

17. Arlt A, Bauer I, Schafmayer C, Tepel J, Muerkoster SS, Brosch M, et al. Increased proteasome subunit protein expression and proteasome activity in colon cancer relate to an enhanced activation of nuclear factor E2-related factor 2 (Nrf2). Oncogene. 2009;28(45):3983–96. Epub 2009/09/08.

18. Chen L, Madura K. Increased proteasome activity, ubiquitin-conjugating enzymes, and eEF1A translation factor detected in breast cancer tissue. Cancer Res. 2005;65(13):5599–606. Epub 2005/07/05.

19. Adams J, Kauffman M. Development of the proteasome inhibitor Velcade (Bortezomib). Cancer Invest. 2004;22(2):304–11. Epub 2004/06/18.

20. Richardson PG, Barlogie B, Berenson J, Singhal S, Jagannath S, Irwin D, et al. A phase 2 study of bortezomib in relapsed, refractory myeloma. N Engl J Med. 2003;348(26):2609–17. Epub 2003/06/27.

21. Caravita T, de Fabritiis P, Palumbo A, Amadori S, Boccadoro M. Bortezomib: efficacy comparisons in solid tumors and hematologic malignancies. Nat Clin Pract Oncol. 2006;3(7):374–87. Epub 2006/07/11.

22. Manasanch EE, Orlowski RZ. Proteasome inhibitors in cancer therapy. Nat Rev Clin Oncol. 2017;14(7):417–33.

23. Voorhees PM, Orlowski RZ. The proteasome and proteasome inhibitors in cancer therapy. Annu Rev Pharmacol Toxicol. 2006;46:189–213. Epub 2006/01/13.

24. Acosta-Alvear D, Cho MY, Wild T, Buchholz TJ, Lerner AG, Simakova O, et al. Paradoxical resistance of multiple myeloma to proteasome inhibitors by decreased levels of 19S proteasomal subunits. eLife. 2015;4:e08153.

25. Tsvetkov P, Mendillo ML, Zhao J, Carette JE, Merrill PH, Cikes D, et al. Compromising the 19S proteasome complex protects cells from reduced flux through the proteasome. eLife. 2015;4.

26. Tsvetkov P, Sokol E, Jin D, Brune Z, Thiru P, Ghandi M, et al. Suppression of 19S proteasome subunits marks emergence of an altered cell state in diverse cancers. Proc Natl Acad Sci U S A. 2017;114(2):382–7.

27. Guo X, Wang X, Wang Z, Banerjee S, Yang J, Huang L, et al. Site-specific proteasome phosphorylation controls cell proliferation and tumorigenesis. Nature cell biology. 2016;18(2):202–12.

28. Tai HC, Besche H, Goldberg AL, Schuman EM. Characterization of the Brain 26S Proteasome and its Interacting Proteins. Frontiers in molecular neuroscience. 2010;3.

29. Myeku N, Clelland CL, Emrani S, Kukushkin NV, Yu WH, Goldberg AL, et al. Tau-driven 26S proteasome impairment and cognitive dysfunction can be prevented early in disease by activating cAMP-PKA signaling. Nature medicine. 2016;22(1):46–53.

30. Livnat-Levanon N, Kevei E, Kleifeld O, Krutauz D, Segref A, Rinaldi T, et al. Reversible 26S proteasome disassembly upon mitochondrial stress. Cell reports. 2014;7(5):1371–80.

31. Tsvetkov P, Myers N, Eliav R, Adamovich Y, Hagai T, Adler J, et al. NADH binds and stabilizes the 26S proteasomes independent of ATP. J Biol Chem. 2014;289(16):11272–81.

32. Lokireddy S, Kukushkin NV, Goldberg AL. cAMP-induced phosphorylation of 26S proteasomes on Rpn6/PSMD11 enhances their activity and the degradation of misfolded proteins. Proc Natl Acad Sci U S A. 2015;112(52):E7176–85.

33. Rousseau A, Bertolotti A. An evolutionarily conserved pathway controls proteasome homeostasis. Nature. 2016.

34. Vilchez D, Boyer L, Morantte I, Lutz M, Merkwirth C, Joyce D, et al. Increased proteasome activity in human embryonic stem cells is regulated by PSMD11. Nature. 2012;489(7415):304–8.

35. Vilchez D, Morantte I, Liu Z, Douglas PM, Merkwirth C, Rodrigues AP, et al. RPN-6 determines C. elegans longevity under proteotoxic stress conditions. Nature. 2012;489(7415):263–8.

36. Caniard A, Ballweg K, Lukas C, Yildirim AO, Eickelberg O, Meiners S. Proteasome function is not impaired in healthy aging of the lung. Aging. 2015;7(10):776–92. Epub 2015/11/06.

37. Walerych D, Lisek K, Sommaggio R, Piazza S, Ciani Y, Dalla E, et al. Proteasome machinery is instrumental in a common gain-of-function program of the p53 missense mutants in cancer. Nature cell biology. 2016;18(8):897–909. Epub 2016/06/28.

38. Kumar MS, Hancock DC, Molina-Arcas M, Steckel M, East P, Diefenbacher M, et al. The GATA2 transcriptional network is requisite for RAS oncogene-driven non-small cell lung cancer. Cell. 2012;149(3):642–55.

39. Luo J, Emanuele MJ, Li D, Creighton CJ, Schlabach MR, Westbrook TF, et al. A genome-wide RNAi screen identifies multiple synthetic lethal interactions with the Ras oncogene. Cell. 2009;137(5):835–48.

40. Petrocca F, Altschuler G, Tan SM, Mendillo ML, Yan H, Jerry DJ, et al. A genome-wide siRNA screen identifies proteasome addiction as a vulnerability of basal-like triple-negative breast cancer cells. Cancer Cell. 2013;24(2):182–96.

41. Hui L, Zheng Y, Yan Y, Bargonetti J, Foster DA. Mutant p53 in MDA-MB-231 breast cancer cells is stabilized by elevated phospholipase D activity and contributes to survival signals generated by phospholipase D. Oncogene. 2006;25(55):7305–10.

42. Inagi R, Ishimoto Y, Nangaku M. Proteostasis in endoplasmic reticulum--new mechanisms in kidney disease. Nat Rev Nephrol. 2014;10(7):369–78.

43. Wang T, Birsoy K, Hughes NW, Krupczak KM, Post Y, Wei JJ, et al. Identification and characterization of essential genes in the human genome. Science. 2015;350(6264):1096–101.

44. Anchoori RK, Karanam B, Peng S, Wang JW, Jiang R, Tanno T, et al. A bis benzylidine piperidone targeting proteasome ubiquitin receptor RPN13/ADRM1 as a therapy for cancer. Cancer Cell. 2013;24(6):791–805.

45. D'Arcy P, Brnjic S, Olofsson MH, Fryknas M, Lindsten K, De Cesare M, et al. Inhibition of proteasome deubiquitinating activity as a new cancer therapy. Nature medicine. 2011;17(12):1636–40.

46. Kemp M. Recent Advances in the Discovery of Deubiquitinating Enzyme Inhibitors. Prog Med Chem. 2016;55:149–92.

47. Deshaies RJ. Proteotoxic crisis, the ubiquitin-proteasome system, and cancer therapy. BMC Biol. 2014;12:94.

48. Mendillo ML, Santagata S, Koeva M, Bell GW, Hu R, Tamimi RM, et al. HSF1 drives a transcriptional program distinct from heat shock to support highly malignant human cancers. Cell. 2012;150(3):549–62.

49. Schop R, Baker A, Chng WJ, et al. Promiscuous mutations activate the noncanonical NF-kappaB pathway in multiple myeloma. Cancer Cell. 2007;12(2):131–44.

50. Nikiforov MA, Riblett M, Tang WH, Gratchouck V, Zhuang D, Fernandez Y, et al. Tumor cell-selective regulation of NOXA by c-MYC in response to proteasome inhibition. Proc Natl Acad Sci U S A. 2007;104(49):19488–93.

51. Tan TT, Degenhardt K, Nelson DA, Beaudoin B, Nieves-Neira W, Bouillet P, et al. Key roles of BIM-driven apoptosis in epithelial tumors and rational chemotherapy. Cancer Cell. 2005;7(3):227–38.

52. Puthalakath H, O'Reilly LA, Gunn P, Lee L, Kelly PN, Huntington ND, et al. ER stress triggers apoptosis by activating BH3-only protein Bim. Cell. 2007;129(7):1337–49.

53. Borkovich KA, Farrelly FW, Finkelstein DB, Taulien J, Lindquist S. hsp82 is an essential protein that is required in higher concentrations for growth of cells at higher temperatures. Mol Cell Biol. 1989;9(9):3919–30.

54. Karras GI, Yi S, Sahni N, Fischer M, Xie J, Vidal M, et al. HSP90 Shapes the Consequences of Human Genetic Variation. Cell. 2017;168(5):856–66 e12.

55. Rutherford SL, Lindquist S. Hsp90 as a capacitor for morphological evolution. Nature. 1998;396(6709):336–42.

56. Taipale M, Jarosz DF, Lindquist S. HSP90 at the hub of protein homeostasis: emerging mechanistic insights. Nat Rev Mol Cell Biol. 2010;11(7):515–28.

57. Nijhawan D, Zack TI, Ren Y, Strickland MR, Lamothe R, Schumacher SE, et al. Cancer vulnerabilities unveiled by genomic loss. Cell. 2012;150(4):842–54.

58. Elsasser S, Schmidt M, Finley D. Characterization of the proteasome using native gel electrophoresis. Methods Enzymol. 2005;398:353–63.

59. Tsvetkov P, Adamovich Y, Elliott E, Shaul Y. E3 ligase STUB1/CHIP regulates NAD(P)H:quinone oxidoreductase 1 (NQO1) accumulation in aged brain, a process impaired in certain Alzheimer disease patients. J Biol Chem. 286(11):8839–45. Epub 2011/01/12.

60. Glickman MH, Rubin DM, Fried VA, Finley D. The regulatory particle of the Saccharomyces cerevisiae proteasome. Mol Cell Biol. 1998;18(6):3149–62.

